# Structure of dimeric full-length human ACE2 in complex with B^0^AT1

**DOI:** 10.1101/2020.02.17.951848

**Authors:** Renhong Yan, Yuanyuan Zhang, Yaning Li, Lu Xia, Qiang Zhou

## Abstract

Angiotensin-converting enzyme 2 (ACE2) is the surface receptor for SARS coronavirus (SARS-CoV), directly interacting with the spike glycoprotein (S protein). ACE2 is also suggested to be the receptor for the new coronavirus (2019-nCoV), which is causing a serious epidemic in China manifested with severe respiratory syndrome. B^0^AT1 (SLC6A19) is a neutral amino acid transporter whose surface expression in intestinal cells requires ACE2. Here we present the 2.9 Å resolution cryo-EM structure of full-length human ACE2 in complex with B^0^AT1. The complex, assembled as a dimer of ACE2-B^0^AT1 heterodimers, exhibits open and closed conformations due to the shifts of the peptidase domains (PDs) of ACE2. A newly resolved Collectrin-like domain (CLD) on ACE2 mediates homo-dimerization. Structural modelling suggests that the ACE2-B^0^AT1 complex can bind two S proteins simultaneously, providing important clues to the molecular basis for coronavirus recognition and infection.

## Introduction

The 2019-nCoV is a type of positive-stranded RNA viruses that causes severe respiratory syndrome in human. The resulting outbreak of the coronavirus disease 2019 (COVID-19) has emerged as a severe epidemic, claiming more than one hundred lives everyday (1, 2). The genome of 2019-nCoV shares ~ 80% identity with that of SARS-CoV and ~ 96% identical to a bat coronavirus (2).

In the case of SARS-CoV, the spike glycoprotein (S protein) on the virion surface mediates receptor recognition and membrane fusion (3, 4). During viral maturation, the trimeric S protein is cleaved into S1 and S2 subunits (4, 5). S1 contains the receptor binding domain (RBD), which directly binds to ACE2 (6), and S2 is responsible for membrane fusion. When S1 binds to the host receptor ACE2, another cleavage site on S2 is exposed and cut by host proteases, a process that is critical for successful viral infection (5, 7, 8). It has been shown that the S protein of 2019-nCoV, same as SARS-CoV, may exploit ACE2 for host infection (2, 9–11). In particular, a most recent *bioRxiv* preprint suggests that the affinity between ACE2 and the RBD of 2019-nCoV is 10-20 times higher than that with the RBD of SARS-CoV (12).

Although ACE2 is known to the public because it is hijacked by coronaviruses, the primary physiological role of ACE2, as its name indicates, is to facilitate maturation of angiotensin, a peptide hormone that controls vasoconstriction and blood pressure. ACE2 is a type I membrane protein expressed in lung, heart, kidney and intestine (13–15). The expression level of ACE2 is associated with cardiovascular diseases (16–18). The full-length ACE2 consists of an N-terminal peptidase domain (PD) and a C-terminal Collectrin-like domain (CLD) that ends with a single transmembrane helix and a ~ 40-residue intracellular segment (13, 19). The PD of ACE2 cleaves angiotensin (Ang) I to produce Ang-(1–9), which is then processed by other enzymes to become Ang-(1–7). ACE2 can also directly process angiotensin II to result in Ang-(1–7) (13, 20).

The PD of ACE2 also provides the direct binding site for the S proteins of coronaviruses. Crystal structure of the claw-like ACE2-PD (21) and structures of the complex between PD and S protein of SARS-CoV have revealed the molecular details of the interaction between the RBD of S protein and PD of ACE2 (6, 22, 23). Apart from PD, the rest of ACE2 is structurally unavailable. The single-TM of ACE2 represents a major challenge for structural determination of the full-length protein.

Recent studies show that ACE2 moonlights as the chaperone for the membrane trafficking of an amino acid transporter, B^0^AT1, also known as SLC6A19 (24). B^0^AT1 mediates uptake of neutral amino acids into intestinal cells in a sodium dependent manner. Its deficiency may cause the Hartnup disorder, an inherited disease with pellagra, cerebellar ataxia and psychosis (25–27). Structures of SLC6 members of *d*DAT (*Drosophila* Dopamine transporter) and human SERT (Serotonin transporter, SLC6A4) have been reported (28, 29). It is unclear how ACE2 interacts with B^0^AT1 and facilitates its membrane trafficking. Our structure of LAT1-4F2hc shows that the cargo and the chaperon interact through both extracellular and transmembrane domains (30). We reasoned that the structure of the full-length ACE2 may be revealed in the presence of B^0^AT1.

Here we report the cryo-EM structure of the full-length human ACE2-B^0^AT1 complex at an overall resolution of 2.9 Å. The complex exists as a dimer of heterodimers. Two conformations of the PD in the ACE2 homodimer, open and close, were extracted from the same dataset. Docking of the structure of the complex between ACE2-PD and the SARS-CoV S protein suggests that two S protein trimers can simultaneously bind to an ACE2 homodimer.

## Structural determination of ACE2-B^0^AT1 complex

The full-length human ACE2 and B^0^AT1, with Strep and FLAG tags on their respective N-terminus, were co-expressed in HEK293F cells and purified through tandem affinity resin and size exclusion chromatography. The complex was eluted in a single mono-disperse peak, indicating high homogeneity (Fig. 1A). Details of cryo-sample preparation, data acquisition, and structural determination are illustrated in Materials and Methods. Eventually, out of 418,140 selected particles, a 3D reconstruction was obtained at an overall resolution of 2.9 Å. An organization of dimer of the ACE2-B^0^AT1 heterodimers was immediately revealed (Fig. 1B). After application of focused refinement and C2 symmetry expansion, the resolution of the extracellular domains was improved to 2.7 Å, while the TM domain remains the same (Figs. 1B, Supplementary Figures 1-3, Table 1).

**Figure 1.**
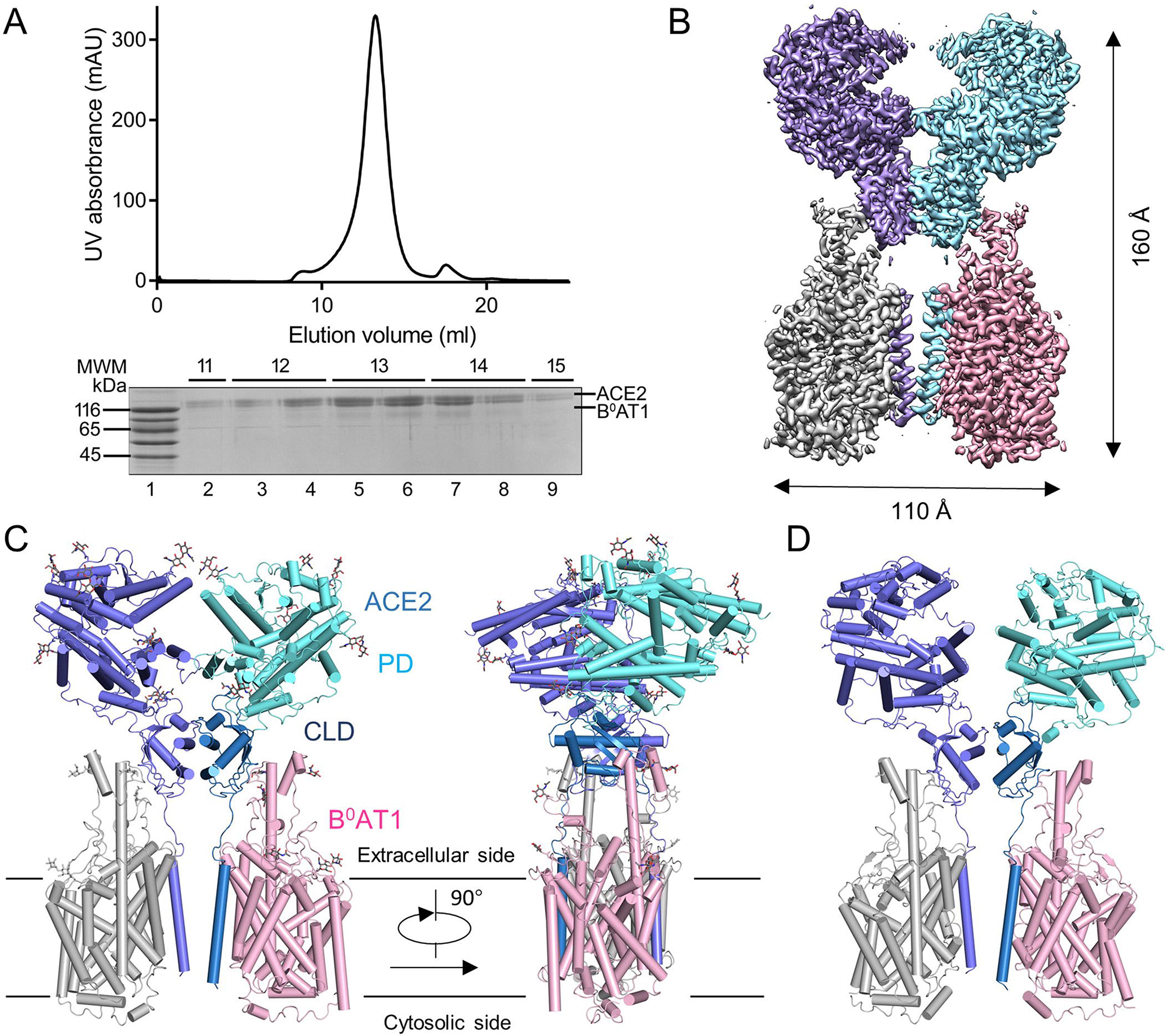
Overall structure of the ACE2-B^0^AT1 complex. **(A)** Representative SEC purification profile of the full-length human ACE2 in complex with B^0^AT1. **(B)** Cryo-EM map of the ACE2-B^0^AT1 complex. The map is generated by merging the focused refined maps shown in Extended Data Figure 2. (**C**) Cartoon representation of the atomic model of the ACE2-B^0^AT1 complex. The glycosylation moieties are shown as sticks. The complex is colored by subunits, with the protease domain (PD) and the Collectrin-like domain (CLD) in one ACE2 protomer colored cyan and blue, respectively. (**D**) An open conformation of the ACE2-B^0^AT1 complex. The two PDs, which contact each other in the “closed” conformation, are separated in the “open” conformation.

The high resolution supported reliable model building. For ACE2, side chains can be assigned to residues 19-768 that contain the PD (residues 19-615) and the newly revealed CLD consisting of a small extracellular domain, a long linker, and the single TM (Fig. 1C). The small domain, sharing a ferredoxin-like fold, will be referred to as the Neck domain because it connects the PD and the TM (Fig. 1C, Supplementary Figure 4). The homo-dimerization is entirely mediated by ACE2, which is sandwiched by B^0^AT1. Both the PD and Neck domain contribute to dimerization, while each B^0^AT1 interacts with the Neck and TM in the adjacent ACE2 (Fig. 1C). The extracellular region is highly glycosylated, with 7 and 5 glycosylation sites on ACE2 and B^0^AT1, respectively.

During classification, another subset with 143,857 particles was processed to an overall resolution of 4.5 Å. It is apparent that while the Neck domain still dimerizes, the PDs are separated from each other in this reconstruction (Fig. 1D, Supplementary Figure 1H-K). We therefore define the two classes as the open and closed conformations. Structural comparison shows that the conformational changes are achieved through rotation of the PD domains only, with the rest of the complex nearly unchanged (Supplementary Movie 1).

## Homodimer interface of ACE2

Dimerization of ACE2 is mainly mediated by the Neck domain, with PD contributing a minor interface (Fig. 2A). To facilitate structural illustrations, we will refer to the two ACE2 protomers A and B, with residues in protomer B labeled with’. Extensive polar interactions are mapped to the interface between N2 and N4 helices on the Neck domain (Fig. 2B). Arg652 and Arg710 in ACE2-A form cation-π interaction with Tyr641’ and Tyr633’ from ACE2-B, respectively. Meanwhile, Arg652 and Arg710 are respectively hydrogen-bonded (H-bond) to Asn638’ and Glu639’, which simultaneously interact with Gln653, in addition to Asn636’. Ser709 and Asp713 from ACE2-A are H-bonded to Arg716’. This extensive network of polar interactions indicates a stable dimer formation.

**Figure 2.**
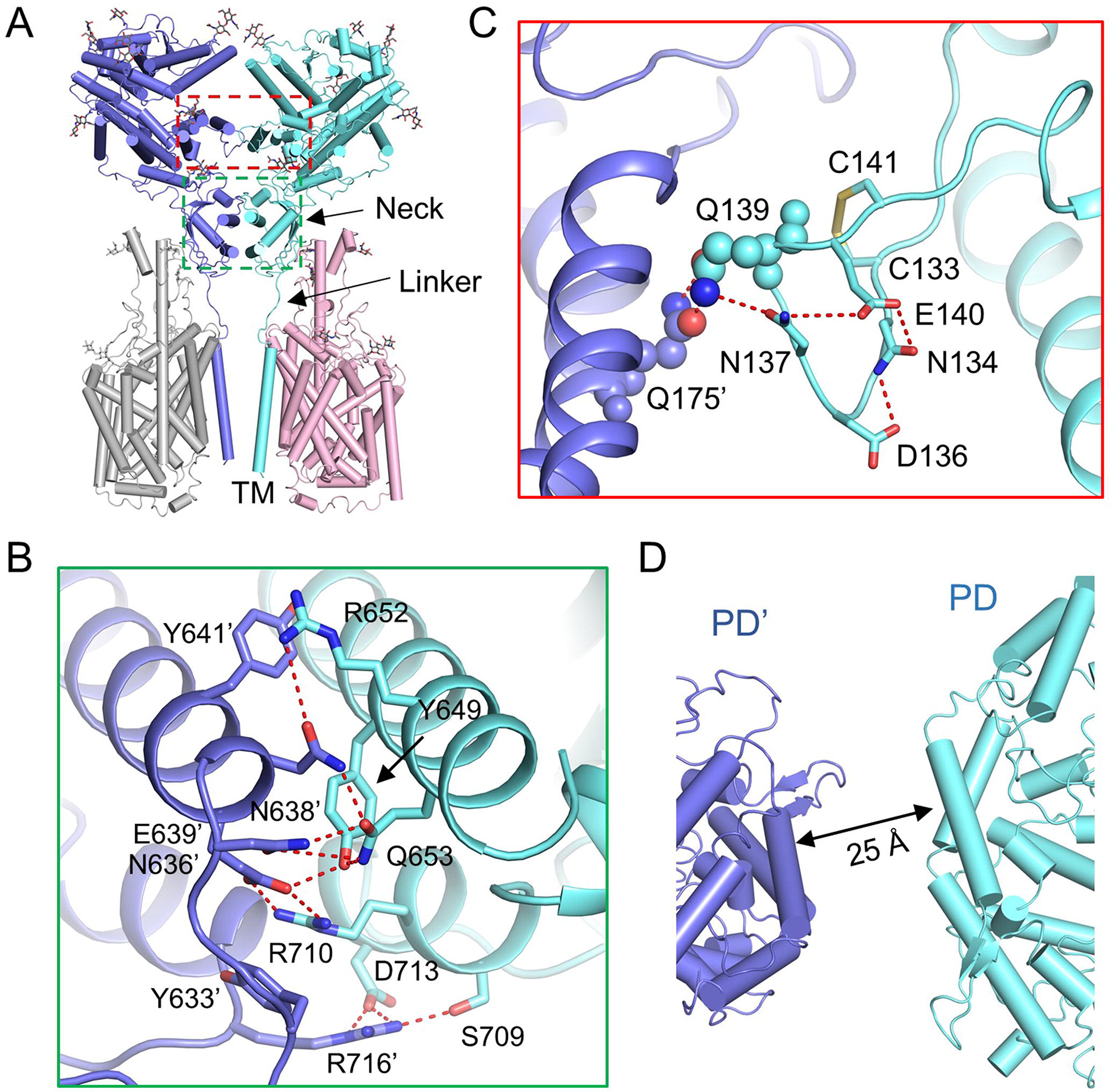
Dimerization interface of ACE2. **(A)** ACE2 dimerizes via two interfaces, the PD and the Neck domain. **(B)** The primary dimeric interface is between the Neck domain in ACE2. Polar interactions are represented by dashed, red lines.**(C)** A weaker interface between PDs of ACE2.**(D)** The PDs no longer contact each other in the open state.

The PD dimer interface appears much weaker with only one pair of interactions between Gln139 and Gln175’ (Fig. 2C). Note that Gln139 is positioned on a loop which is stabilized by a disulfide bond between Cys133 and Cys141 as well as multiple intro-loop polar interactions (Fig. 2C). The weak interaction is consistent with the presence of the open conformation, in which the interface between the Neck domains remains the same while the PDs are separated from each other by ~ 25 Å (Fig. 2D, Supplementary Movie 1).

## Two trimeric S proteins can bind to an ACE2 dimer simultaneously

The available structures for the complex between the SARS-CoV S protein and ACE2 are limited to the isolated PD only. We next examined whether a dimeric ACE2 can simultaneously accommodate two S protein trimers. When the S protein-PD complex (PDB code: 6ACJ) is superimposed with the PD in the closed or open states of the ACE2-B^0^AT1 complex, the PD domains can be aligned with root mean squared deviation of ~ 1.9 Å over ~ 550 pairs of C_α_ atoms with the closed conformation or of ~ 4.9 Å over ~ 590 pairs of C_α_ atoms with the open conformation (Fig. 3A, B, Supplementary Figure 5). In both states, S protein is positioned on the outer side of ACE2 dimer. Therefore, a dimeric ACE2 can accommodate two S protein trimers, simultaneously. This observation immediately implies potential clustering between dimeric ACE2 and trimeric S proteins, which may be important for invagination of the membrane and endocytosis of the viral particle, a process similar to other receptor-mediated endocytosis.

**Figure 3.**
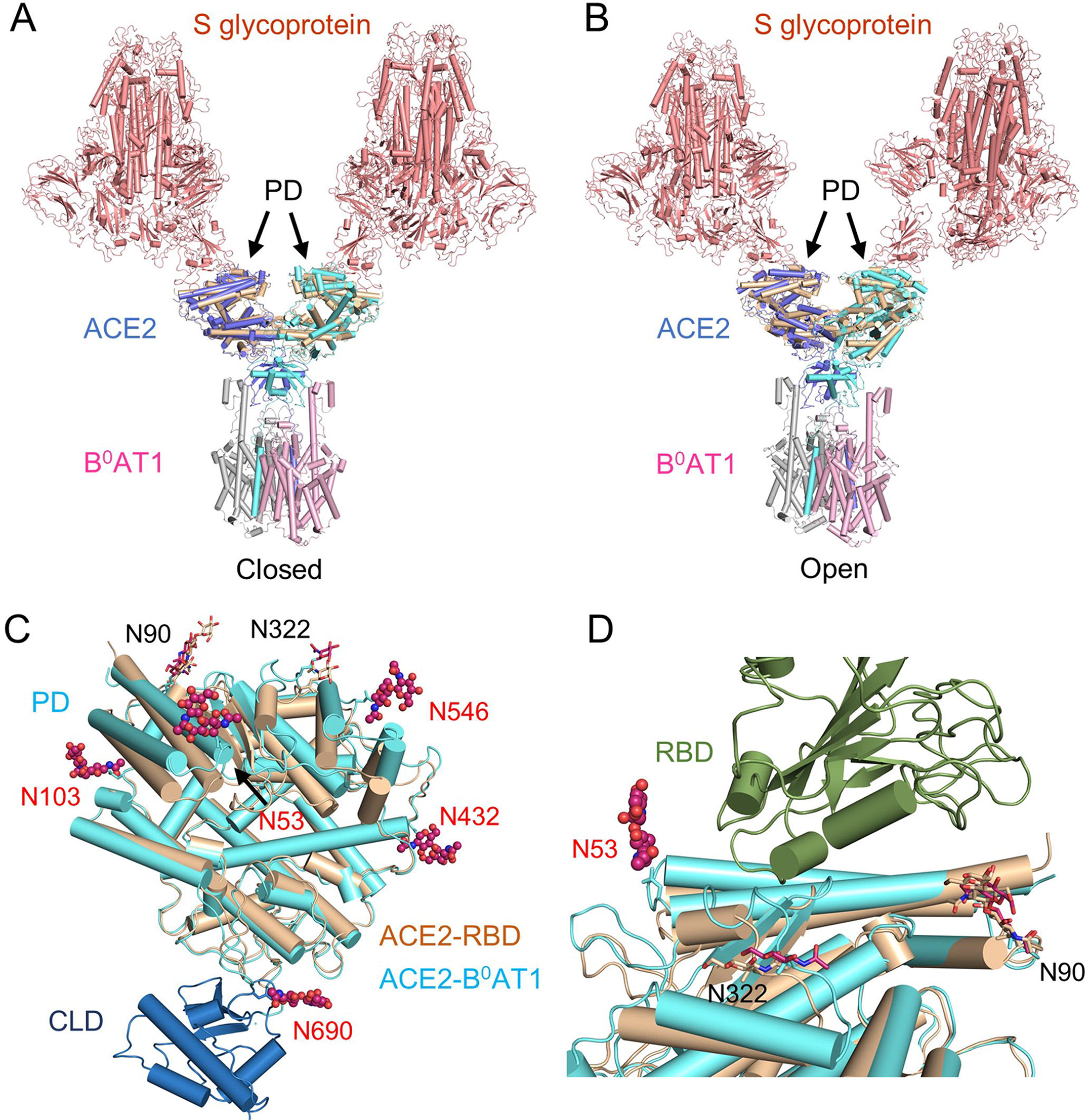
Two trimeric Spike glycoprotein of SARS-CoV can bind to a dimeric ACE2 simultaneously. **(A)** Structure comparison between the ACE2-B^0^AT1 complex and the complex of the spike glycoprotein (S protein) of SARS-CoV and PD. The structure of the S protein-PD complex (PDB ID: 6ACJ) is superimposed to the PD of the closed ACE2-B^0^AT1 complex.**(B)** is same to **(A)**, but with the open conformation.**(C)** Glycosylation sites in ACE2. In total seven glycosylation sites are identified in the structure of the ACE2-B^0^AT1 complex. Two are the same with previous report. The ones that are found in this study are shown as spheres and labeled red.**(D)** Sugar moieties may participate in S protein binding. Structure of the complex between ACE2-PD and the receptor binding domain (RBD) of the S protein of SARS-CoV (PDB ID: 2AJF) is overlaid with the ACE2-B^0^AT1 complex. The binding site for RBD is surrounded by three glycosylation sites on PD.

Because our proteins were expressed in mammalian cells, whereas the published ACE2-PD was obtained from baculovirus-infected insect cells, more glycosylation sites, six vs two, were observed on the surface of PD (Fig. 3C). The significance of glycosylation in the recognition between viruses and receptors has drawn increasing attention. Among the observed glycosylation sites on PD, three are in the vicinity of the RBD binding site (Fig. 3D). Two, Asn90 and Asn322, were observed previously (6), and one, Asn53, is revealed in this study. It has been shown that chloroquine, which can interfere with the terminal glycosylation of ACE2 (31), inhibits SARS-CoV infection. It remains to be shown whether these sugar moieties directly participate in S protein binding.

## The interfaces between ACE2 and B^0^AT1

As the PD and Neck domain are connected to the TM though an elongated linker, it is likely that the extended conformation is stabilized by B^0^AT1. Indeed, a close look at the heterodimer shows extensive interface along one side of the Neck domain and TM of ACE2 with an extended TM7 and TM4 of B^0^AT1 (Fig. 4A), reminiscent to the interactions between 4F2hc and LAT1 (30, 32).

**Figure 4.**
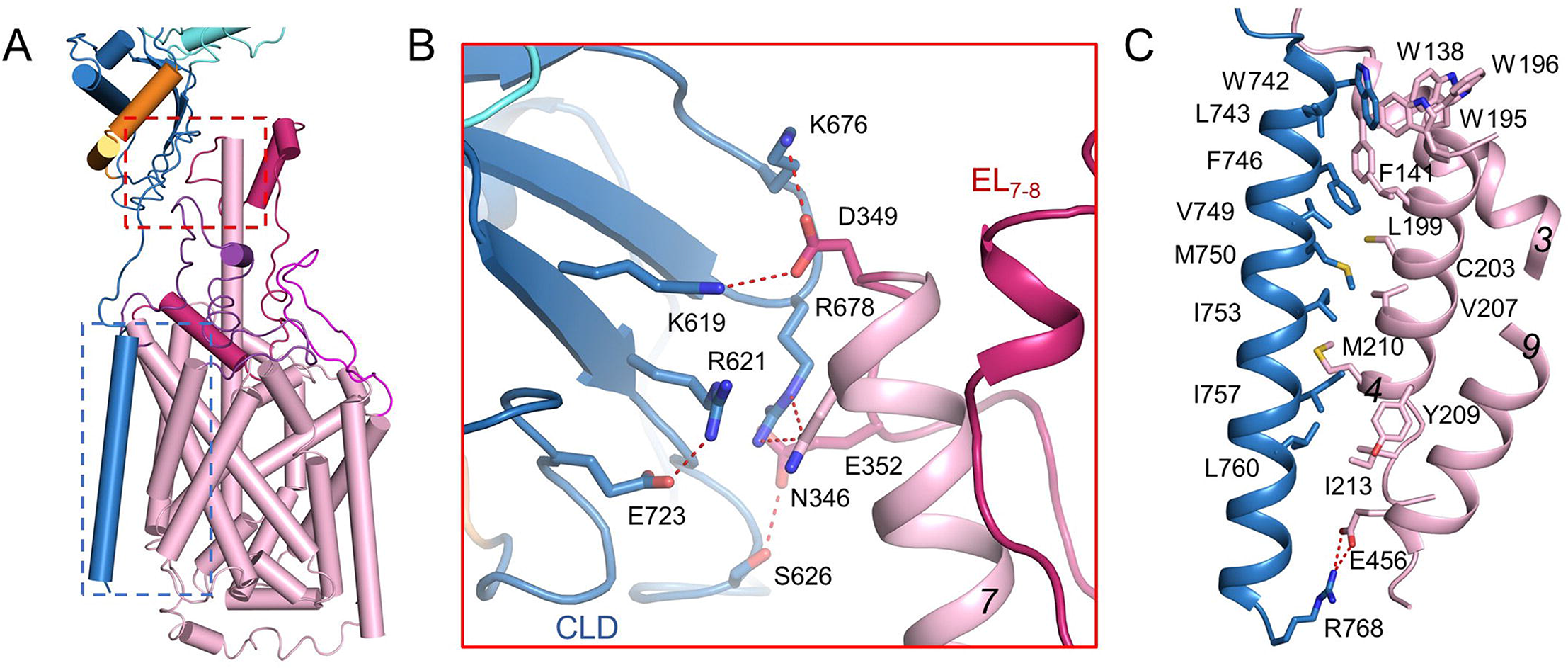
Interactions of ACE2 and B^0^AT1. **(A)** ACE2 interacts with B^0^AT1 both through extracellular domains and within membrane. Only one ACE2-B^0^AT1 heterodimer is shown for clarity. The coloured boxes correspond to those analysed in details in panels **B** and **C**. **(B)** The extracellular interface 1 between PD and the extended TM7 of B^0^AT1. **(C)** The interactions between the TM helix of ACE2 and TMs 3, 4, and 9 of B^0^AT1.

Compared to dDAT and other LeuT-fold transporters, TM7 of B^0^AT1 is particularly long with its C-terminus extruding out of the membrane by ~ 25 Å (Supplementary Figure 6A). The extended TM7 segment becomes an organizing center to coordinate other extracellular segments of B^0^AT1 that participate in the complex formation with ACE2. On the extracellular side, the Neck domain of ACE2 associates with the C-terminal end of TM7 and the ensuing sequence of EL_7-8_ (extracellular loop between TMs 7 and 8) of B^0^AT1 via a number of polar or charged residues. Asn346 and Glu352 of B^0^AT1 bind to Ser626 and Arg678 of ACE2, respectively. Asp349 of B^0^AT1 interacts with Lys676 of ACE2 directly and with Lys619 and Arg621 through a water molecule (Fig. 4B).

Below the Neck domain, the linker is close to the EL_3-4_ loop of B^0^AT1 (Supplementary Figure 6B). However, the density for this region is relatively weak. No specific interactions can be identified. Further down to the membrane, the TM of ACE2 closely packs against TM3 and TM4 of B^0^AT1 via van der Waals contacts throughout the height of the lipid bilayer. At the C-terminal cytoplasmic tip of the TM, Arg768 of ACE2 interacts with Glu456 on TM9 of B^0^AT1 to further secure the binding of the two proteins (Fig. 4C).

## Discussion

Although ACE2 is a chaperon for B^0^AT1, our focus is on ACE2 in this study. With the stabilization by B^0^AT1, we elucidated the structure of the full-length ACE2, which provides important insight into the molecular basis for viral infection by SARS-CoV as well as 2019-nCoV.

There has been no report on the oligomerization state of ACE2. Our structure reveals the assembly of a dimer of heterodimers between ACE2 and B^0^AT1. B^0^AT1 is not involved in homo-dimerization, whereas the contacts between the Neck domains of ACE2 are extensive. Therefore ACE2 likely exists as a homodimer even in the absence of B^0^AT1. This analysis provides immediate clue to the molecular basis for viral infection by SARS-CoV or 2019-nCoV.

Cleavage of the S protein of SARS-CoV was reported to be facilitated by cathepsin L in endosomes, indicating a mechanism of receptor-mediated endocytosis (8). The discovery of ACE2 as a dimer and the compatibility of simultaneous binding of dimeric ACE2 with trimeric S protein suggests possible clustering that may facilitate membrane invagination for endocytosis. Further characterizations are required to examine the interactions between ACE2 and the viral particle as well as the effect of co-factors such as B^0^AT1 and integrin on this process.

For membrane fusion, cleavage of the C-terminal segment, especially residues 697 to 716 (Supplementary Figure 4), of ACE2 by proteases, such as transmembrane protease serine 2 (TMPRSS2), can enhance S protein-driven viral entry (33, 34). In our present structure, residues of 697-716 form helix N3 and N4 in the Neck domain and map to the dimeric interface of ACE2. Presence of B^0^AT1 may block the access of TMPRSS2 to the cutting site on ACE2. The expression distribution of ACE2 is broader than B^0^AT1. In addition to kidney and intestine, where B^0^AT1 is primarily expressed, ACE2 is expressed in lung and heart (26). It remains to be tested whether B^0^AT1 can suppress SARS-CoV infection by blocking ACE2 cleavage.

We started this work because of our serious concern with the COVID-19. Therefore the focus of structural analysis is mainly on ACE2. In fact, B^0^AT1, as an important nutrient importer, exhibits many interesting structural features that are different from other LeuT-fold transporters. We will aim for structural determination of B^0^AT1 in more conformations and present structural analysis of B^0^AT1 in separate studies.

In summary, our structural reveals the high-resolution structures of full-length ACE2 in a dimeric assembly. Modelling analysis suggests simultaneous binding of two S protein trimer of coronavirus to an ACE2 dimer. The structure presents here serves as an important framework to dissect the mechanism of 2019-nCoV infection and may facilitate development of potential therapeutics.

## Acknowledgments

We thank the Cryo-EM Facility and Supercomputer Center of Westlake University for providing cryo-EM and computation support, respectively. This work was funded by the National Natural Science Foundation of China (projects 31971123, 81920108015, 31930059) and the Key R&D Program of Zhejiang Province (2020C04001).

## Author contributions

Q.Z. and R.Y. conceived the project. Q.Z and R.Y. designed the experiments. All authors did the experiments. Q.Z. R.Y., Y.L. and Y.Z. contributed to data analysis. Q.Z. and R.Y. wrote the manuscript.

## References

1. Zhu N, Zhang D, Wang W, Li X, Yang B, Song J, Zhao X, Huang B, Shi W, Lu R, Niu P, Zhan F, Ma X, Wang D, Xu W, Wu G, Gao GF, Tan W, China Novel Coronavirus I, Research T. A Novel Coronavirus from Patients with Pneumonia in China, 2019. N Engl J Med. 2020. doi: 10.1056/NEJMoa2001017. PubMed PMID: 31978945.

2. Zhou P, Yang XL, Wang XG, Hu B, Zhang L, Zhang W, Si HR, Zhu Y, Li B, Huang CL, Chen HD, Chen J, Luo Y, Guo H, Jiang RD, Liu MQ, Chen Y, Shen XR, Wang X, Zheng XS, Zhao K, Chen QJ, Deng F, Liu LL, Yan B, Zhan FX, Wang YY, Xiao GF, Shi ZL. A pneumonia outbreak associated with a new coronavirus of probable bat origin. Nature. 2020. Epub 2020/02/06. doi: 10.1038/s41586-020-2012-7. PubMed PMID: 32015507.

3. Gallagher TM, Buchmeier MJ. Coronavirus spike proteins in viral entry and pathogenesis. Virology. 2001;279(2):371–4. doi: 10.1006/viro.2000.0757. PubMed PMID: 11162792.

4. Simmons G, Zmora P, Gierer S, Heurich A, Pohlmann S. Proteolytic activation of the SARS-coronavirus spike protein: cutting enzymes at the cutting edge of antiviral research. Antiviral Res. 2013;100(3):605–14. doi: 10.1016/j.antiviral.2013.09.028. PubMed PMID: 24121034; PMCID: PMC3889862.

5. Belouzard S, Chu VC, Whittaker GR. Activation of the SARS coronavirus spike protein via sequential proteolytic cleavage at two distinct sites. Proc Natl Acad Sci U S A. 2009;106(14):5871–6. doi: 10.1073/pnas.0809524106. PubMed PMID: 19321428; PMCID: PMC2660061.

6. Li F, Li W, Farzan M, Harrison SC. Structure of SARS coronavirus spike receptor-binding domain complexed with receptor. Science. 2005;309(5742):1864–8. doi: 10.1126/science.1116480. PubMed PMID: 16166518.

7. Millet JK, Whittaker GR. Host cell proteases: Critical determinants of coronavirus tropism and pathogenesis. Virus Res. 2015;202:120–34. doi: 10.1016/j.virusres.2014.11.021. PubMed PMID: 25445340; PMCID: PMC4465284.

8. Simmons G, Gosalia DN, Rennekamp AJ, Reeves JD, Diamond SL, Bates P. Inhibitors of cathepsin L prevent severe acute respiratory syndrome coronavirus entry. Proc Natl Acad Sci U S A. 2005;102(33):11876–81. doi: 10.1073/pnas.0505577102. PubMed PMID: 16081529; PMCID: PMC1188015.

9. Li W, Moore MJ, Vasilieva N, Sui J, Wong SK, Berne MA, Somasundaran M, Sullivan JL, Luzuriaga K, Greenough TC, Choe H, Farzan M. Angiotensin-converting enzyme 2 is a functional receptor for the SARS coronavirus. Nature. 2003;426(6965):450–4. doi: 10.1038/nature02145. PubMed PMID: 14647384.

10. Kuba K, Imai Y, Rao S, Gao H, Guo F, Guan B, Huan Y, Yang P, Zhang Y, Deng W, Bao L, Zhang B, Liu G, Wang Z, Chappell M, Liu Y, Zheng D, Leibbrandt A, Wada T, Slutsky AS, Liu D, Qin C, Jiang C, Penninger JM. A crucial role of angiotensin converting enzyme 2 (ACE2) in SARS coronavirus-induced lung injury. Nat Med. 2005;11(8):875–9. doi: 10.1038/nm1267. PubMed PMID: 16007097.

11. Hoffmann M, Kleine-Weber H, Krüger N, Müller M, Drosten C, Pöhlmann S. The novel coronavirus 2019 (2019-nCoV) uses the SARS-coronavirus receptor ACE2 and the cellular protease TMPRSS2 for entry into target cells. bioRxiv. 2020.

12. Wrapp D, Wang N, Corbett KS, Goldsmith JA, Hsieh C-L, Abiona O, Graham BS, McLellan JS. Cryo-EM Structure of the 2019-nCoV Spike in the Prefusion Conformation. bioRxiv. 2020.

13. Donoghue M, Hsieh F, Baronas E, Godbout K, Gosselin M, Stagliano N, Donovan M, Woolf B, Robison K, Jeyaseelan R, Breitbart RE, Acton S. A novel angiotensin-converting enzyme-related carboxypeptidase (ACE2) converts angiotensin I to angiotensin 1-9. Circ Res. 2000;87(5):E1–9. doi: 10.1161/01.res.87.5.e1. PubMed PMID: 10969042.

14. Zhang H, Kang Z, Gong H, Xu D, Wang J, Li Z, Cui X, Xiao J, Meng T, Zhou W, Liu J, Xu H. The digestive system is a potential route of 2019-nCov infection: a bioinformatics analysis based on single-cell transcriptomes. bioRxiv. 2020.

15. Zhao Y, Zhao Z, Wang Y, Zhou Y, Ma Y, Zuo W. Single-cell RNA expression profiling of ACE2, the putative receptor of Wuhan 2019-nCov. bioRxiv. 2020.

16. Crackower MA, Sarao R, Oudit GY, Yagil C, Kozieradzki I, Scanga SE, Oliveira-dos-Santos AJ, da Costa J, Zhang L, Pei Y, Scholey J, Ferrario CM, Manoukian AS, Chappell MC, Backx PH, Yagil Y, Penninger JM. Angiotensin-converting enzyme 2 is an essential regulator of heart function. Nature. 2002;417(6891):822–8. doi: 10.1038/nature00786. PubMed PMID: 12075344.

17. Raizada MK, Ferreira AJ. ACE2: a new target for cardiovascular disease therapeutics. J Cardiovasc Pharmacol. 2007;50(2):112–9. doi: 10.1097/FJC.0b013e3180986219. PubMed PMID: 17703127.

18. Zisman LS, Keller RS, Weaver B, Lin Q, Speth R, Bristow MR, Canver CC. Increased angiotensin-(1-7)-forming activity in failing human heart ventricles: evidence for upregulation of the angiotensin-converting enzyme Homologue ACE2. Circulation. 2003;108(14):1707–12. doi: 10.1161/01.CIR.0000094734.67990.99. PubMed PMID: 14504186.

19. Zhang H, Wada J, Hida K, Tsuchiyama Y, Hiragushi K, Shikata K, Wang H, Lin S, Kanwar YS, Makino H. Collectrin, a collecting duct-specific transmembrane glycoprotein, is a novel homolog of ACE2 and is developmentally regulated in embryonic kidneys. J Biol Chem. 2001;276(20):17132–9. doi: 10.1074/jbc.M006723200. PubMed PMID: 11278314.

20. Hamming I, Cooper ME, Haagmans BL, Hooper NM, Korstanje R, Osterhaus AD, Timens W, Turner AJ, Navis G, van Goor H. The emerging role of ACE2 in physiology and disease. J Pathol. 2007;212(1):1–11. doi: 10.1002/path.2162. PubMed PMID: 17464936.

21. Towler P, Staker B, Prasad SG, Menon S, Tang J, Parsons T, Ryan D, Fisher M, Williams D, Dales NA, Patane MA, Pantoliano MW. ACE2 X-ray structures reveal a large hinge-bending motion important for inhibitor binding and catalysis. J Biol Chem. 2004;279(17):17996–8007. doi: 10.1074/jbc.M311191200. PubMed PMID: 14754895.

22. Song W, Gui M, Wang X, Xiang Y. Cryo-EM structure of the SARS coronavirus spike glycoprotein in complex with its host cell receptor ACE2. PLoS Pathog. 2018;14(8):e1007236. doi: 10.1371/journal.ppat.1007236. PubMed PMID: 30102747; PMCID: PMC6107290.

23. Kirchdoerfer RN, Wang N, Pallesen J, Wrapp D, Turner HL, Cottrell CA, Corbett KS, Graham BS, McLellan JS, Ward AB. Receptor binding and proteolysis do not induce large conformational changes in the SARS-CoV spike. bioRxiv. 2018.

24. Kowalczuk S, Broer A, Tietze N, Vanslambrouck JM, Rasko JE, Broer S. A protein complex in the brush-border membrane explains a Hartnup disorder allele. FASEB J. 2008;22(8):2880–7. doi: 10.1096/fj.08-107300. PubMed PMID: 18424768.

25. Seow HF, Broer S, Broer A, Bailey CG, Potter SJ, Cavanaugh JA, Rasko JE. Hartnup disorder is caused by mutations in the gene encoding the neutral amino acid transporter SLC6A19. Nat Genet. 2004;36(9):1003–7. doi: 10.1038/ng1406. PubMed PMID: 15286788.

26. Kleta R, Romeo E, Ristic Z, Ohura T, Stuart C, Arcos-Burgos M, Dave MH, Wagner CA, Camargo SR, Inoue S, Matsuura N, Helip-Wooley A, Bockenhauer D, Warth R, Bernardini I, Visser G, Eggermann T, Lee P, Chairoungdua A, Jutabha P, Babu E, Nilwarangkoon S, Anzai N, Kanai Y, Verrey F, Gahl WA, Koizumi A. Mutations in SLC6A19, encoding B^0^AT1, cause Hartnup disorder. Nat Genet. 2004;36(9):999–1002. doi: 10.1038/ng1405. PubMed PMID: 15286787.

27. Broer A, Klingel K, Kowalczuk S, Rasko JE, Cavanaugh J, Broer S. Molecular cloning of mouse amino acid transport system B0, a neutral amino acid transporter related to Hartnup disorder. J Biol Chem. 2004;279(23):24467–76. doi: 10.1074/jbc.M400904200. PubMed PMID: 15044460.

28. Penmatsa A, Wang KH, Gouaux E. X-ray structure of dopamine transporter elucidates antidepressant mechanism. Nature. 2013;503(7474):85–90. doi: 10.1038/nature12533. PubMed PMID: 24037379; PMCID: PMC3904663.

29. Coleman JA, Green EM, Gouaux E. X-ray structures and mechanism of the human serotonin transporter. Nature. 2016;532(7599):334–9. doi: 10.1038/nature17629. PubMed PMID: 27049939; PMCID: PMC4898786.

30. Yan R, Zhao X, Lei J, Zhou Q. Structure of the human LAT1-4F2hc heteromeric amino acid transporter complex. Nature. 2019. doi: 10.1038/s41586-019-1011-z. PubMed PMID: 30867591.

31. Vincent MJ, Bergeron E, Benjannet S, Erickson BR, Rollin PE, Ksiazek TG, Seidah NG, Nichol ST. Chloroquine is a potent inhibitor of SARS coronavirus infection and spread. Virol J. 2005;2:69. doi: 10.1186/1743-422X-2-69. PubMed PMID: 16115318; PMCID: PMC1232869.

32. Lee Y, Wiriyasermkul P, Jin C, Quan L, Ohgaki R, Okuda S, Kusakizako T, Nishizawa T, Oda K, Ishitani R, Yokoyama T, Nakane T, Shirouzu M, Endou H, Nagamori S, Kanai Y, Nureki O. Cryo-EM structure of the human L-type amino acid transporter 1 in complex with glycoprotein CD98hc. Nat Struct Mol Biol. 2019;26(6):510–7. doi: 10.1038/s41594-019-0237-7. PubMed PMID: 31160781.

33. Shulla A, Heald-Sargent T, Subramanya G, Zhao J, Perlman S, Gallagher T. A transmembrane serine protease is linked to the severe acute respiratory syndrome coronavirus receptor and activates virus entry. J Virol. 2011;85(2):873–82. doi: 10.1128/JVI.02062-10. PubMed PMID: 21068237; PMCID: PMC3020023.

34. Heurich A, Hofmann-Winkler H, Gierer S, Liepold T, Jahn O, Pohlmann S. TMPRSS2 and ADAM17 cleave ACE2 differentially and only proteolysis by TMPRSS2 augments entry driven by the severe acute respiratory syndrome coronavirus spike protein. J Virol. 2014;88(2):1293–307. doi: 10.1128/JVI.02202-13. PubMed PMID: 24227843; PMCID: PMC3911672.

