## Supplementary for "Structure of dimeric full-length human ACE2 in complex with B^0^AT1"

### Methods

#### Protein preparation

The cDNAs for full-length human $B^0AT1$ (accession number: NM_001003841) and ACE2 (accession number: NM_001371415) were subcloned into pCAG respectively. An N-terminal FLAG tag was fused to $B^0AT1$, and one Strep tag was fused after the N-terminal signal peptide of ACE2 using a standard two-step PCR.

HEK 293F cells (Invitrogen) were cultured in SMM 293T-II medium (Sino Biological Inc.) at 37 °C under 5% $CO_2$ in a Multitron-Pro shaker (Infors, 130 rpm). To co-express $B^0AT1$ and ACE2, the cells were transiently transfected with the plasmids and polyethylenimines (PEIs) (Polysciences) when the cell density reached approximately 2.0 $\times 10^6$/ml. For transfection one liter of cell culture, about 0.75 mg plasmids for $B^0AT1$ and 0.75 mg plasmids for ACE2 were premixed with 3 mg PEIs in 50 ml of fresh medium for 15 mins before adding to cell culture. The transfected cells were cultured for 48-60 hours before harvesting.

For purification of the $B^0AT1$ and ACE2 complex, the cells were collected in a buffer containing 25 mM Tris, pH 8.0, 150 mM NaCl, and three protease inhibitors , aprotinin (1.3 μg/ml, AMRESCO), pepstatin (0.7 μg/ml, AMRESCO), and leupeptin (5 μg/ml, AMRESCO). The membrane fraction was solubilized at 4 °C for 2 hours with 1% (w/v) glyco diosgenin (GDN, Anatrace) and the cell debris was removed by centrifugation at 18,700 g for 45 mins. The supernatant was loaded to anti-FLAG M2 affinity resin (Sigma). After rinsed with the wash buffer containing 25 mM Tris, pH 8.0, 150 mM NaCl, and 0.02% GDN (w/v), the protein was eluted with wash buffer plus 0.2 mg/ml FLAG peptide. The eluent was further purified by Strep-Tactin Sepharose (IBA). After eluted with the wash buffer supplemented with 4 mM desthiobiotin (IBA), the eluent was then concentrated and subject to size-exclusion chromatography (Superose 6 Increase 10/300 GL, GE Healthcare) in the buffer containing 25 mM Tris, pH 8.0, 150 mM NaCl, and 0.02% GDN. The peak fractions were collected and concentrated for EM analysis.

**Cryo-EM sample preparation and data acquisition**

For cryo-sample preparation, the purified protein complex was concentrated to ~ 8 mg/ml. Aliquots (3.3 µl) of the protein complex were placed on glow-discharged holey carbon grids (Quantifoil Au R1.2/1.3), which were blotted for 3.0 s or 3.5 s and flash-frozen in liquid ethane cooled by liquid nitrogen with Vitrobot (Mark IV, Thermo Fisher Scientific). The cryo grids were transferred to a Titan Krios operating at 300 kV equipped with Gatan K3 Summit detector and GIF Quantum energy filter. Movie stacks were automatically collected using AutoEMation (35), with a slit width of 20 eV on the energy filter and a defocus range from -1.2 µm to -2.2 µm in super-resolution mode at a nominal magnification of 81,000×. Each stack was exposed for 2.56 s with an exposure time of 0.08 s per frame, resulting in a total of 32 frames per stack. The total dose rate was approximately 50 $e^-/Å^2$ for each stack. The stacks were motion corrected with MotionCor2 (36) and binned 2-fold, resulting in a pixel size of 1.087 Å/pixel. Meanwhile, dose weighting was performed (37). The defocus values were estimated with Gctf (38).

#### Data processing

Particles were automatically picked using Relion 3.0.6 (39-42) from manually selected micrographs. After 2D classification with Relion, good particles were selected and subject to 3D classification with Relion with C2 symmetry against an initial model generated with Relion (the ACE2-B$^0$AT1 complex). The open and closed conformation particles were selected and subject to local defocus correction (38), 3D auto-refinement and post-processing. To improve the map quality of the closed conformation, the dataset was further refined with adapted mask applied on the extracellular domains and the TM domains, respectively. For TM domains, the dataset was symmetry-expanded before refinement. For the dataset of the open conformation of the ACE2-B$^0$AT1 complex, the dataset was further subject to 2 rounds of heterogeneous refinement and non-uniform refinement with cryoSPARC (43).

The resolution was estimated with the gold-standard Fourier shell correlation 0.143 criterion (44) with high-resolution noise substitution (45). Refer to Supplemental Figures S1-S2 and Supplemental Table S1 for details of data collection and processing.

#### Model building and structure refinement

Model building of the ACE2-B$^0$AT1 complex was performed by MDFF (46) of the published structure (PDB ID: 6ACJ) for the PD domain of ACE2 or *ab initio* with Phenix (47) and Coot (48) for the other parts based on the focused-refined cryo-EM maps with aromatic residues as landmarks, most of which were clearly visible in the cryo-EM map. Each residue was manually checked with the chemical properties taken into consideration during model building. Several segments, whose corresponding densities were invisible, were not modeled.

Structural refinement was performed in Phenix with secondary structure and geometry restraints to prevent overfitting. To monitor the potential overfitting, the model was refined against one of the two independent half maps from the gold-standard 3D refinement approach. Then, the refined model was tested against the other map. Statistics associated with data collection, 3D reconstruction and model building is summarized in Supplementary Table S1.

**Supplementary Table S1 | Data collection, 3D reconstruction and model statistic**

| | | |
|---|---|---|
| **Data collection** | | |
| EM equipment | Titan Krios (Thermo Fisher Scientific) | |
| Voltage (kV) | 300 | |
| Detector | Gatan K3 Summit | |
| Energy filter | Gatan GIF Quantum, 20 eV slit | |
| Pixel size (Å) | 1.087 | |
| Electron dose (e-/Å2) | 50 | |
| Defocus range (μm) | -1.2 ~ -2.2 | |
| Number of collected micrographs | 3,952 | |
| Number of selected micrographs | 3,904 | |
| Conformation | closed | open |
| **3D Reconstruction** | | |
| Software | Relion 3.0 | cryoSPARC |
| Number of used particles | 418,140 | 143,857 |
| Resolution (Å) | 2.9 | 4.5 |
| Symmetry | C2 | C1 |
| Map sharpening B factor (Å2) | -90 | -140 |
| **Refinement** | | |
| Software | Phenix | |
| Cell dimensions | | |
| a=b=c (Å) | 313.056 | |
| α=β=γ (˚) | 90 | |
| Model composition | | |
| Protein residues | 2,708 | 2,708 |
| Side chains assigned | 2,708 | 2,708 |
| Sugar | 38 | |
| ligand | 4 | |
| Phospholipid | 4 | |
| Water | 30 | |
| R.m.s deviations | | |
| Bonds length (Å) | 0.010 | 0.005 |
| Bonds Angle (˚) | 1.085 | 1.123 |
| Ramachandran plot statistics (%) | | |
| Preferred | 91.11 | 90.55 |
| Allowed | 8.74 | 9.30 |
| Outlier | 0.15 | 0.15 |

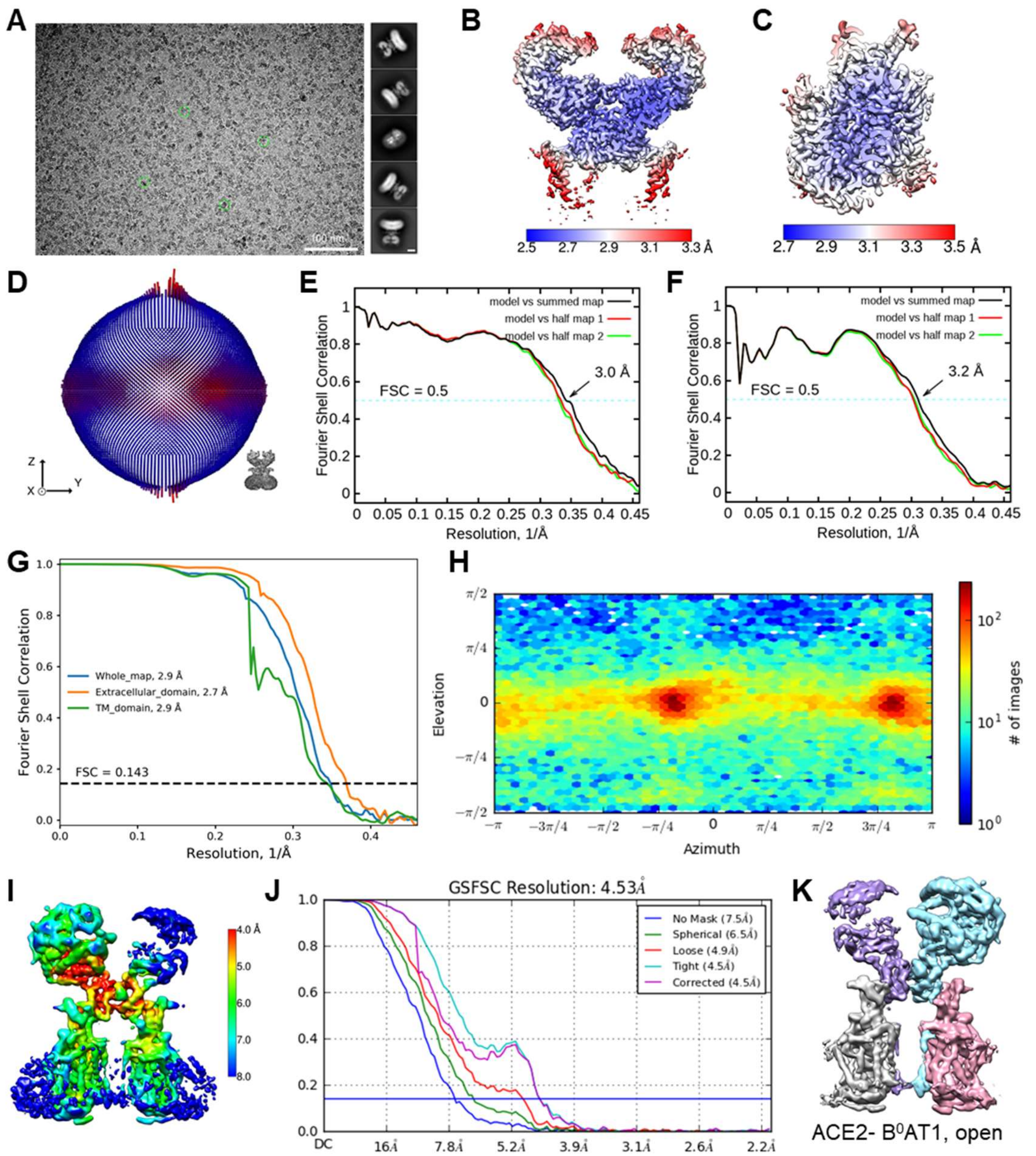


**Supplementary Fig. S1. Cryo-EM analysis of ACE2- B$^0$AT1 complex in closed and open conformation.** **(A)** Representative electron micrograph and 2D class averages of cryo-EM particle images. The scale bar in 2D class averages represents 10 nm. **(B)** and **(C)** Local resolution map for the 3D reconstruction of extracellular region and TM region of closed ACE2- B$^0$AT1 complex. **(D)** Euler angle distribution of closed ACE2- B$^0$AT1 complex in the final 3D reconstruction. **(E)** FSC curve of the refined model of closed ACE2- B$^0$AT1 versus the extracellular domain that it is refined against (black); of the model refined against the first half map versus the same map (red); and of the model refined against the first half map versus the second half map (green). The small difference between the red and green curves indicates that the refinement of the atomic coordinates did not suffer from overfitting. **(F)** FSC curve of the refined model of TM domain of closed ACE2- $B^0$AT1 complex, which is the same as the (E). **(G)** Gold standard FSC curve of the overall structure (blue), extracellular domain (orange) and TM domain (green) of closed ACE2- $B^0$AT1 complex. **(H)** Euler angle distribution of open ACE2- $B^0$AT1 complex in the final 3D reconstruction in cryoSPARC. **(I)** Local resolution map for the 3D reconstruction of open ACE2- $B^0$AT1 complex. **(J)** Gold standard FSC curve of open ACE2- $B^0$AT1 complex is estimated by cryoSPARC. **(K)** Cryo-EM map of the ACE2- $B^0$AT1 complex in the open conformation. The map is coloured by subunits.

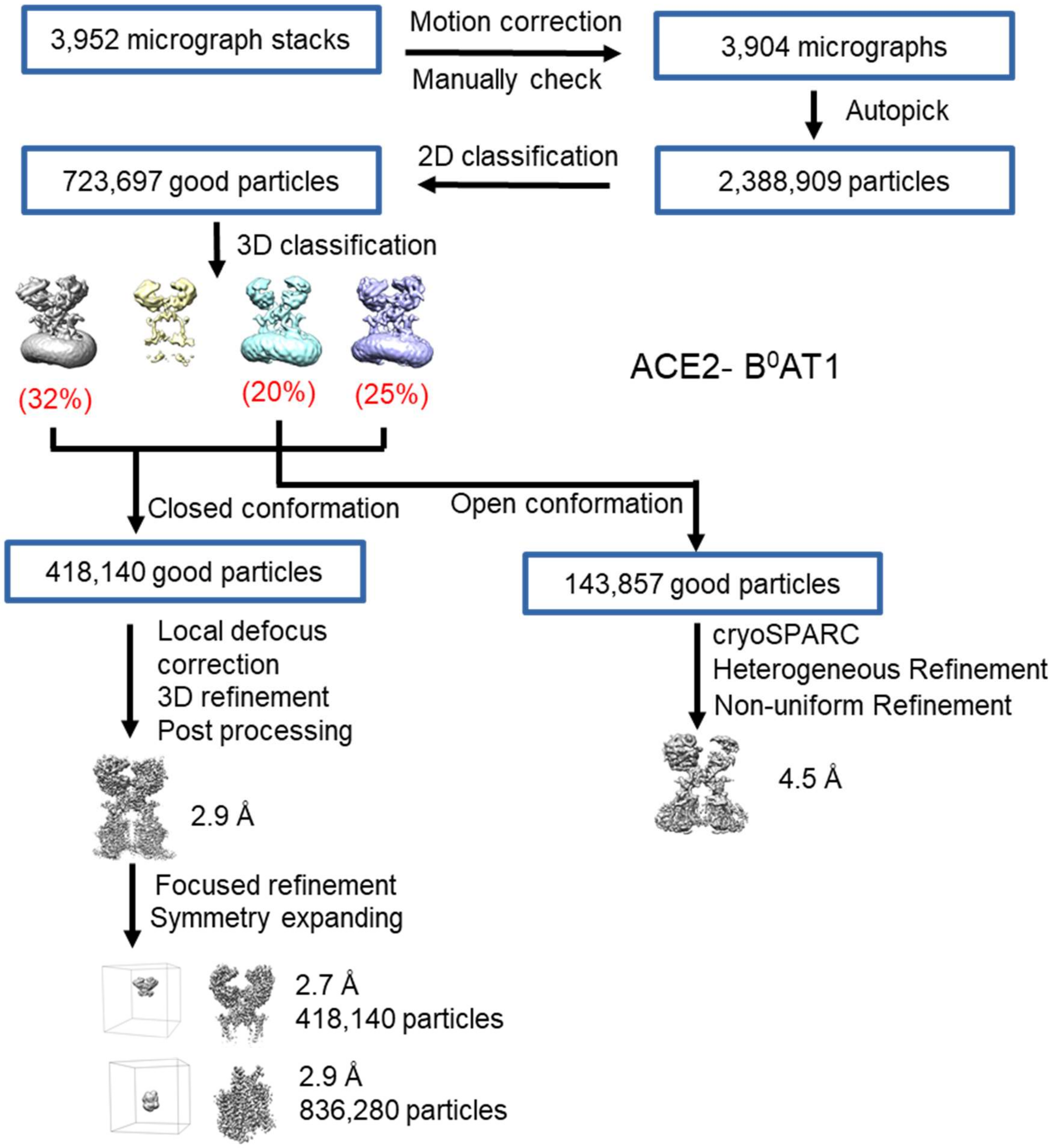


**Supplementary Fig. S2. Flowchart for cryo-EM data processing.** Please see the "Data Processing" section in Methods for details.

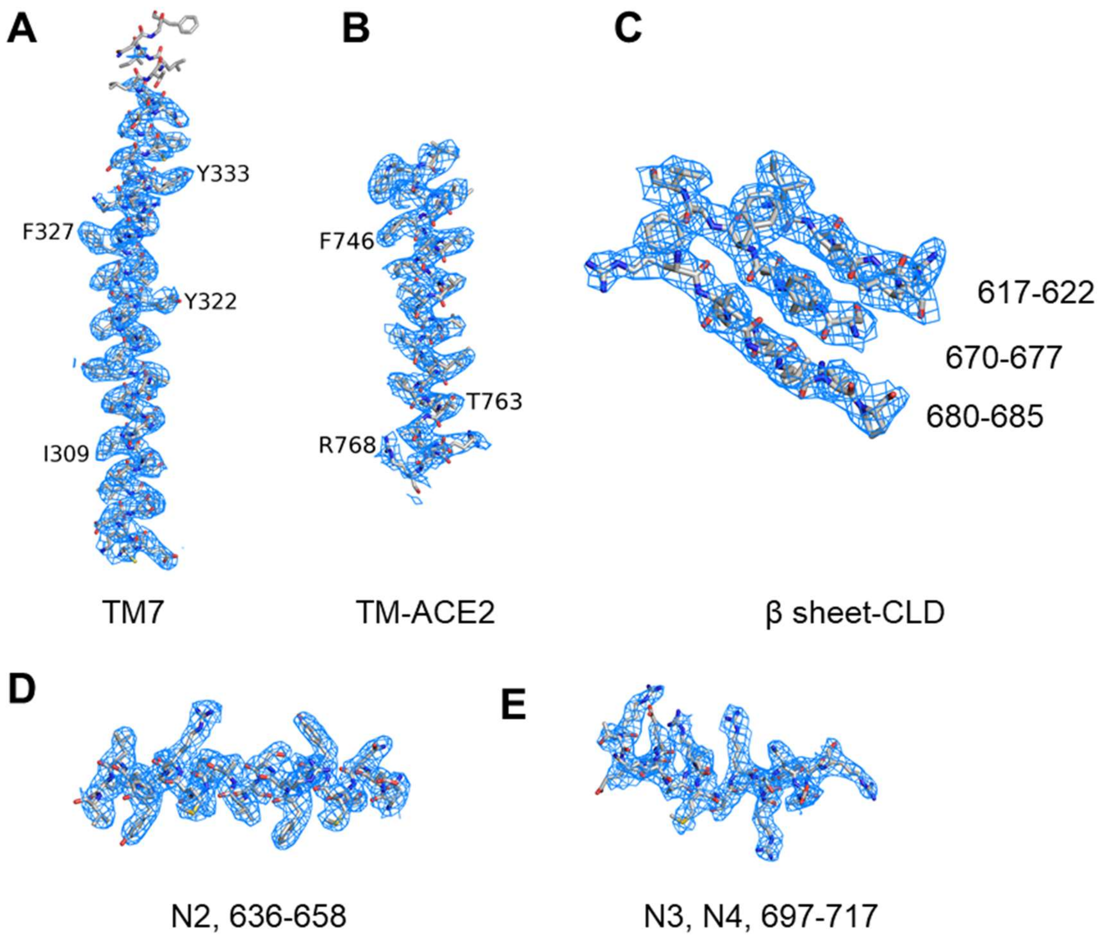


**Supplementary Fig. S3. Representative Cryo-EM densities.** Shown here are the cryo-EM maps of indicated segments of ACE2-$B^0$AT1 complex in the “closed” conformation. All densities are generated in PyMOL and contoured at 8 σ.

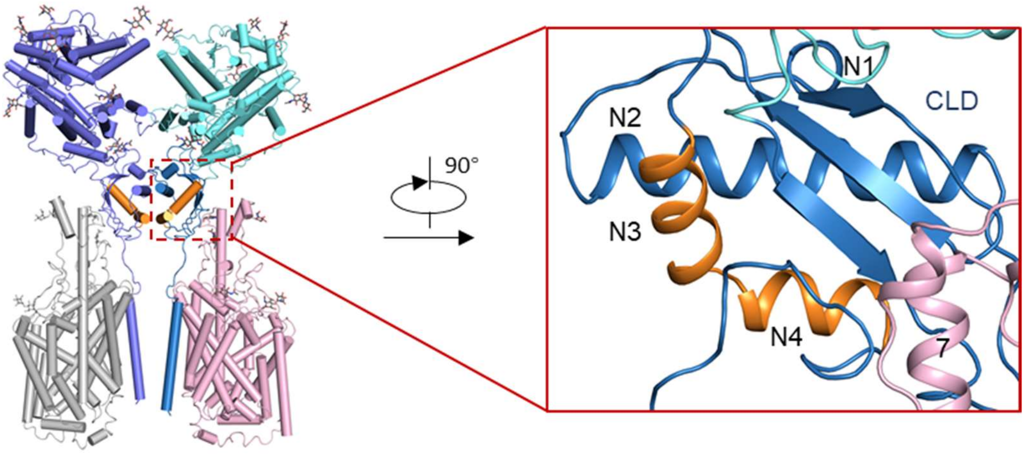


**Supplementary Fig. S4. Structure of the Neck domain in CLD.** The Neck domain exhibits a ferredoxin-like fold. The four α-helices in CLD are labelled N1-N4. The segment containing the predicted cleavage site is colored orange.

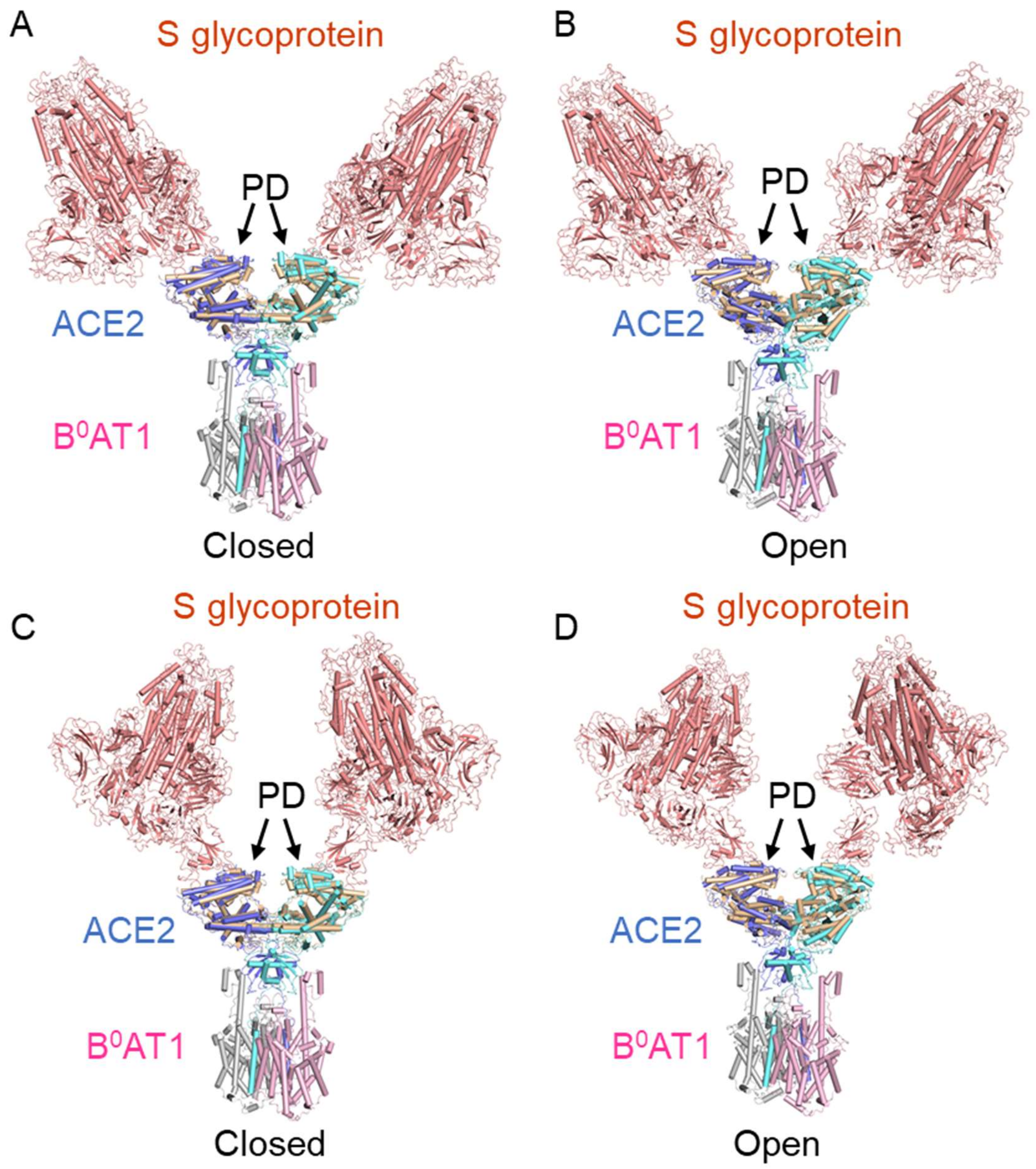


**Supplementary Fig. S5. Structural alignment of the ACE2-B$^0$AT1 complex with PD of ACE2 in complex with S glycoprotein of SARS-CoV.** **(A)** and **(B)** Alignment of the closed and open conformation of ACE2-B$^0$AT1 complex with PD of ACE2 in complex with S glycoprotein of SARS-CoV (PDB ID: 6ACG), respectively. **(C)** and **(D)** are same to (A) and (B), respectively, but using PD of ACE2 in complex with S glycoprotein of SARS-CoV with PDB code 6ACK.

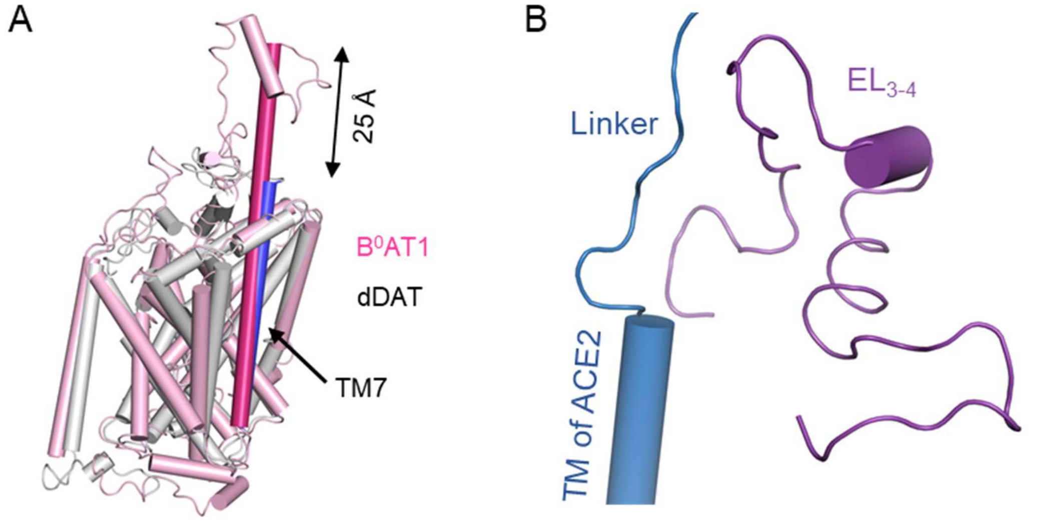


**Supplementary Fig. S6. $B^0$AT1 has a particularly long TM7.** **(A)** $B^0$AT1 possesses an extended TM7. Shown here is the structural comparison between $B^0$AT1 (pink) and dDAT (grey, PDB ID:4XPF). The TM7 segment in $B^0$AT1 and dDAT are highlighted in magenta and blue, respectively. **(B)** The linker of CLD domain is close to the $EL_{3-4}$ loop of $B^0$AT1.

**Supplementary Movie S1.** The structural morph between the closed and open conformation of the ACE2-$B^0AT1$ complex.

**Supplementary References**

35. Lei J, Frank J. Automated acquisition of cryo-electron micrographs for single particle reconstruction on an FEI Tecnai electron microscope. Journal of structural biology. 2005;150(1):69-80. Epub 2005/03/31. doi: 10.1016/j.jsb.2005.01.002. PubMed PMID: 15797731.

36. Zheng SQ, Palovcak E, Armache JP, Verba KA, Cheng Y, Agard DA. MotionCor2: anisotropic correction of beam-induced motion for improved cryo-electron microscopy. Nature methods. 2017;14(4):331-2. Epub 2017/03/03. doi: 10.1038/nmeth.4193. PubMed PMID: 28250466; PMCID: PMC5494038.

37. Grant T, Grigorieff N. Measuring the optimal exposure for single particle cryo-EM using a 2.6 A reconstruction of rotavirus VP6. eLife. 2015;4:e06980. Epub 2015/05/30. doi: 10.7554/eLife.06980. PubMed PMID: 26023829; PMCID: PMC4471936.

38. Zhang K. Gctf: Real-time CTF determination and correction. Journal of structural biology. 2016;193(1):1-12. Epub 2015/11/26. doi: 10.1016/j.jsb.2015.11.003. PubMed PMID: 26592709; PMCID: PMC4711343.

39. Zivanov J, Nakane T, Forsberg BO, Kimanius D, Hagen WJ, Lindahl E, Scheres SH. New tools for automated high-resolution cryo-EM structure determination in RELION-3. eLife. 2018;7. Epub 2018/11/10. doi: 10.7554/eLife.42166. PubMed PMID: 30412051; PMCID: PMC6250425.

40. Kimanius D, Forsberg BO, Scheres SH, Lindahl E. Accelerated cryo-EM structure determination with parallelisation using GPUs in RELION-2. eLife. 2016;5. doi: 10.7554/eLife.18722. PubMed PMID: 27845625.

41. Scheres SH. RELION: implementation of a Bayesian approach to cryo-EM structure determination. Journal of structural biology. 2012;180(3):519-30. Epub 2012/09/25. doi: 10.1016/j.jsb.2012.09.006. PubMed PMID: 23000701; PMCID: PMC3690530.

42. Scheres SH. A Bayesian view on cryo-EM structure determination. Journal of molecular biology. 2012;415(2):406-18. Epub 2011/11/22. doi: 10.1016/j.jmb.2011.11.010. PubMed PMID: 22100448; PMCID: PMC3314964.

43. Punjani A, Rubinstein JL, Fleet DJ, Brubaker MA. cryoSPARC: algorithms for rapid unsupervised cryo-EM structure determination. Nature methods. 2017;14(3):290-6. Epub 2017/02/07. doi: 10.1038/nmeth.4169. PubMed PMID: 28165473.

44. Rosenthal PB, Henderson R. Optimal determination of particle orientation, absolute hand, and contrast loss in single-particle electron cryomicroscopy. Journal of molecular biology. 2003;333(4):721-45. Epub 2003/10/22. PubMed PMID: 14568533.

45. Chen S, McMullan G, Faruqi AR, Murshudov GN, Short JM, Scheres SH, Henderson R. High-resolution noise substitution to measure overfitting and validate resolution in 3D structure determination by single particle electron cryomicroscopy. Ultramicroscopy. 2013;135:24-35. Epub 2013/07/23. doi: 10.1016/j.ultramic.2013.06.004. PubMed PMID: 23872039; PMCID: PMC3834153.

46. Trabuco LG, Villa E, Mitra K, Frank J, Schulten K. Flexible fitting of atomic structures into electron microscopy maps using molecular dynamics. Structure (London, England : 1993). 2008;16(5):673-83. Epub 2008/05/09. doi: 10.1016/j.str.2008.03.005. PubMed PMID: 18462672; PMCID: PMC2430731.

47. Adams PD, Afonine PV, Bunkoczi G, Chen VB, Davis IW, Echols N, Headd JJ, Hung LW, Kapral GJ, Grosse-Kunstleve RW, McCoy AJ, Moriarty NW, Oeffner R, Read RJ, Richardson DC, Richardson JS, Terwilliger TC, Zwart PH. PHENIX: a comprehensive Python-based system for macromolecular structure solution. Acta crystallographica Section D, Biological crystallography. 2010;66(Pt 2):213-21. Epub 2010/02/04. doi: 10.1107/S0907444909052925. PubMed PMID: 20124702; PMCID: 2815670.

48. Emsley P, Lohkamp B, Scott WG, Cowtan K. Features and development of Coot. Acta crystallographica Section D, Biological crystallography. 2010;66(Pt 4):486-501. Epub 2010/04/13. doi: 10.1107/S0907444910007493. PubMed PMID: 20383002; PMCID: 2852313.
